# Using selection by non-antibiotic stressors to sensitize bacteria to antibiotics

**DOI:** 10.1101/628834

**Authors:** Jeff Maltas, Brian Krasnick, Kevin B. Wood

**Affiliations:** Department of Biophysics, University of Michigan, Ann Arbor, USA; Department of Physics, University of Michigan, Ann Arbor, USA

## Abstract

Bacterial resistance to one antibiotic is frequently accompanied by crossresistance to other drugs. Similarly, non-antibiotic selective forces, from biocides to osmotic stress, have been shown to decrease antibiotic susceptibility, often the result of shared, non-specific resistance mechanisms. On the other hand, evolved resistance to particular antibiotics may also be associated with increased *sensitivity* to other drugs, highlighting evolutionary constraints that could form the basis for novel anti-resistance strategies. While recent studies indicate this collateral sensitivity is common between antibiotics, much less is known about potentially sensitizing effects of non-antibiotic stressors. In this study, we use laboratory evolution to investigate adaptation of *E. faecalis*, an opportunistic bacterial pathogen, to a broad collection of environmental agents, ranging from antibiotics and biocides to extreme pH and osmotic stress. We find that non-antibiotic selection frequently leads to increased sensitivity to other conditions, including multiple antibiotics. Using population sequencing and whole genome sequencing of single isolates from the evolved populations, we identify multiple mutations in genes previously linked with resistance to the selecting conditions, including genes corresponding to known drug targets or multi-drug efflux systems previously tied to collateral sensitivity. Finally, we hypothesized based on the measured sensitivity profiles that sequential rounds of antibiotic and non-antibiotic selection may lead to hypersensitive populations by harnessing the orthogonal collateral effects of particular pairs of selective forces. To test this hypothesis, we show experimentally that populations evolved to a sequence of linezolid (an oxazolidinone antibiotic) and sodium benzoate (a common preservative) exhibit increased sensitivity to more stressors than adaptation to either condition alone. The results demonstrate how sequential adaptation to drug and non-drug environments can be used to sensitize bacterial to antibiotics and highlight new potential strategies for exploiting shared constraints governing adaptation to diverse environmental challenges.

## INTRODUCTION

The emergence of drug resistance is continually shrinking an ever-smaller pool of drugs necessary for the successful treatment of infectious disease and cancer (1, 2,3,4,5,6,7). The evolution of resistance is a complex stochastic process that may depend on spatiotemporal dynamics of the host environment (8, 9,10,11,12,13,14). In addition, resistance evolution in fluctuating or multi-agent environments may be driven by phenotypic trade-offs reflecting conflicting evolutionary goals. For example, recent studies have shown that acquiring resistance to a single antibiotic frequently leads to a change in the susceptibility to a different antibiotic, a phenomenon known as collateral sensitivity or cross resistance (15, 16, 17, 18, 19, 20, 21, 22, 23, 24, 25, 26). While the molecular mechanisms of collateral sensitivity have been identified in several specific cases-for example, modulation of proton-motor force underlies increased sensitivity to some antibiotics induced in aminoglycoside-resistant mutants (18)-they are generally difficult to uncover and may vary by species and drug, making them an ongoing focus of research. At the same time, a number of recent studies have shown that systems-level approaches based on phenotypic profiling may help identify statistical properties of these collateral effects, even when molecular mechanisms are not fully known (19,16,17, 24, 27, 28, 26).

In addition to antibiotics, many studies have shown that exposure to non-antibiotic conditions, such as heavy metals, biocides, extreme temperatures, acidic or osmotic stress, and even growth media may also lead to reduced susceptibility to antimicrobials (29, 30, 31, 32, 33, 34, 35, 36, 37, 38). For example, adaptation to the antiseptic chlorhexidine (CHX) was recently shown to be associated with collateral resistance to daptomycin, a lipopeptide antibiotic used to treat multidrug-resistant Gram-positive infections (33). On the other hand, antibiotic resistant strains may exhibit increased sensitivity to antimicrobial peptides (39), and bacteria undergoing long-term evolution without drug generally show decreased antibiotic resistance (40). As a whole, these studies point to overlapping evolutionary constraints that govern adaptation to a large and chemically diverse collection of deleterious environments. In turn, they raise the question of whether non-antibiotic stressors-which are frequently encountered in both clinical and natural environments-might play an important role in the evolution of drug resistance and, at the same time, represent an untapped set of environmental “levers” for steering evolutionary trajectories.

While there has been extensive progress identifying the molecular mechanisms governing cross-resistance between specific pairs of antibiotic and non-antibiotic stressors, relatively little is known about the systems-level properties of these evolutionary trade-offs. Does adaptation to non-antibiotic stressors frequently lead to modulated antibiotic resistance, or are these effects relatively rare, restricted-perhaps-to structurally or mechanistically similar agents? When these collateral effects appear, are they dominated by cross-resistance, pointing to an ever-accelerating march to resistant pathogens with broad multi-agent resistance? Or do these conditions co-select for increased sensitivities, potentially leading to multi-agent environmental sequences that trap cells in evolutionarily vulnerable states? Recent approaches that leverage similar incompatible evolutionary objectives have revolutionized our view of multidrug therapies (41). Non-antibiotic stressors may offer a complementary set of unappreciated selective forces for simultaneously sensitizing pathogens to multiple drugs.

In this work we start to answer some of these questions using laboratory evolution and phenotypic profiling in an opportunistic bacterial pathogen. Specifically, we investigate phenotypic collateral effects arising during bacterial adaptation to 6 antibiotics and 7 non-antibiotic environments, including common biocides, extreme pH, and osmotic stress. As a model system, we focus on *E. faecalis*, a Gram-positive bacterial species frequently found in the gastrointestinal tracts of humans. *E. faecalis* can survive in a range of harsh environments, making it a good candidate for adaptation to many different environmental conditions. At the same time, *E. faecalis* is an important clinical pathogen that contributes to multiple human infections, including urinary tract infections and infective endocarditis (42, 43, 44, 45).

In a recent study, we used laboratory evolution to characterize the phenotypic collateral sensitivity profiles between multiple antibiotics in *E. faecalis* (26). In this study, we show that collateral resistance and sensitivity are also surprisingly common between more general environmental stressors, both between different non-antibiotic stressors and between antibiotics and non-antibiotic conditions. While the specific resistance profiles vary between independent populations, even when selected by the same condition, the collateral sensitivities remain common. For example, 25 of 32 isolates selected by the antimicrobial triclosan (TCS) exhibited increased sensitivity to at least one of the 6 antibiotics tested. Finally, we show experimentally that populations evolved to a sequence of two conditions (the antibiotic linezolid and the preservative sodium benzoate (NaBz)) can induce increased sensitivity to more conditions than adaptation to either stressor alone. The results demonstrate how sequential adaptation to drug and non-drug environments can be used to sensitize bacterial to antibiotics and highlight new potential approaches for leveraging evolutionary trade-offs inherent in adaptation to diverse environments.

## RESULTS

### Collateral effects between antibiotic and non-antibiotic stressors are common

To investigate collateral effects between antibiotic and non-antibiotic conditions, we exposed populations of *E. faecalis* strain V583 to increasing concentrations of a single condition for up to 60 days (approximately 450 generations) via serial passage evolution (Figure 1A, Methods). We repeated this laboratory evolution for 7 different (non-antibiotic) selecting conditions, including extreme pH, osmotic stress, biocides, and preservatives (Table 2). Following laboratory evolution, we isolated a single colony (“mutant”) from each population and measured its susceptibility to all 7 conditions as well as to 6 antibiotics spanning multiple classes (Table 2) via high-throughput dose-response experiments. In addition, we measured susceptibility of 6 previously isolated strains (one for each antibiotic; strains were originally isolated in (26)) to all 7 non-antibiotic stressors. To quantify resistance to each condition, we estimated the half maximal inhibitory concentration (IC_50_) for all 13 isolates, as well as isolates from the ancestral populations, to each of the 13 conditions (Methods; Figure 1B). For each isolate-condition combination, we then calculate *c* ≡ log_2_ (IC_50,*Mut*_/IC_50,*WT*_), the log-scaled fold change in IC_50_ of the mutant (relative to ancestral strains) (Figure 1C). Resistance therefore corresponds to *c* > 0 and sensitivity to *c* < 0.

**FIG 1.**
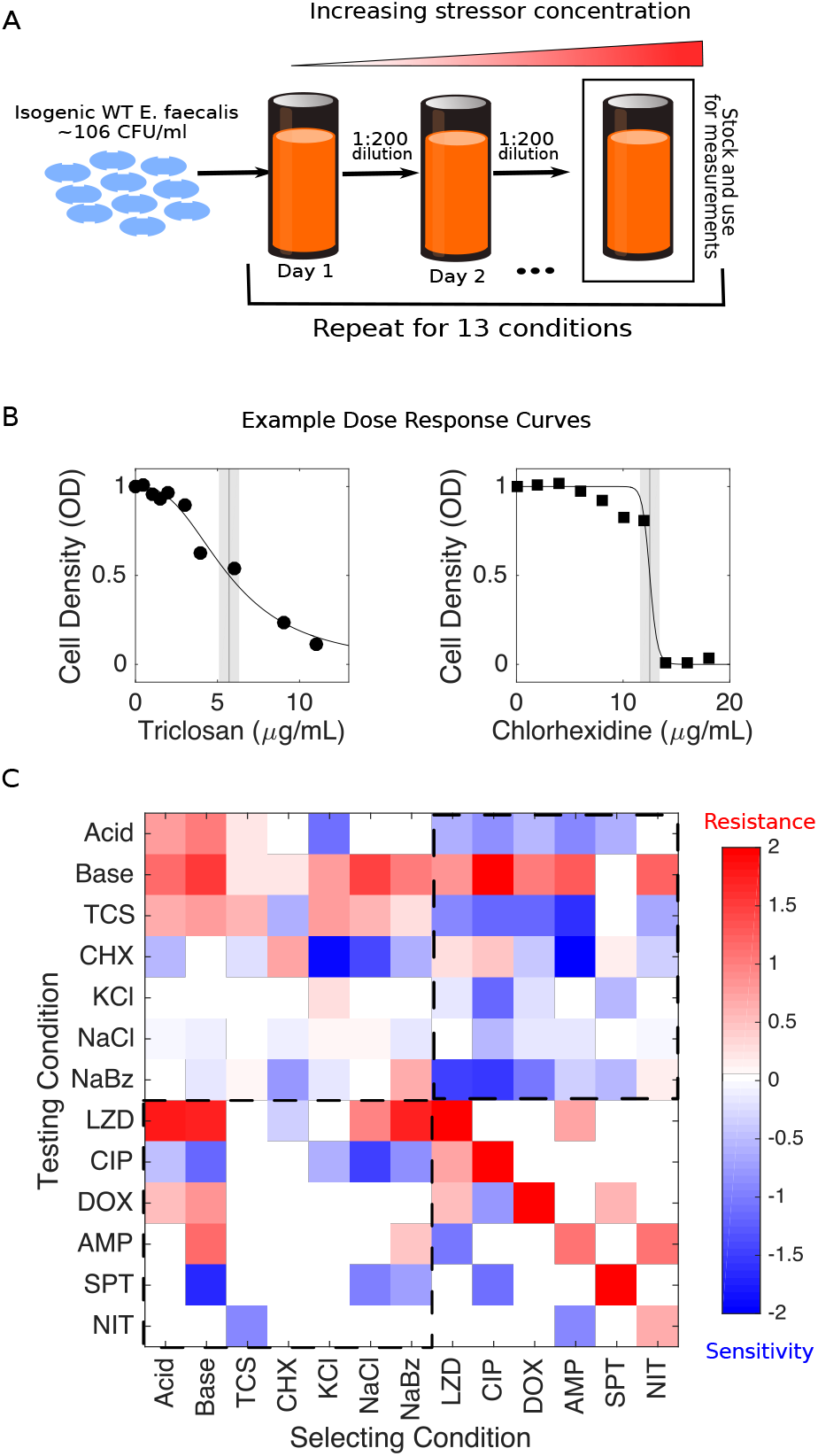
Laboratory evolution reveals collateral sensitivity and cross resistance between antibiotics and environmental stressors in *E. faecalis*. A. Populations of *E. faecalis* V583 were exposed to increasing concentrations of a single selecting condition over multiple days via serial passage experiments (Methods). The evolution was repeated for 13 different selecting conditions, including six antibiotics and seven non-antibiotic stressors (Table I). At the end of the evolution experiment, a single colony was isolated from each population and tested for modulated sensitivity to each of the 13 environmental conditions. B. Example dose response curves for isolates selected by TCS (left) and chlorhexidine (right). Vertical line represents estimated half-maximal inhibitory concentration (IC_50_), with shaded regions confidence intervals (95%). C. Resistance (red) or sensitivity (blue) to each condition is quantified using the (log_2_-transformed) fold change in the IC_50_ for the selected isolate relative to that of ancestral (V583) cells. Dashed regions correspond to antibiotic susceptibilities of non-antibiotic selected isolates (lower left) and, conversely, non-antibiotic susceptibilities of antibiotic selected isolates (upper right).

**TABLE 1.**
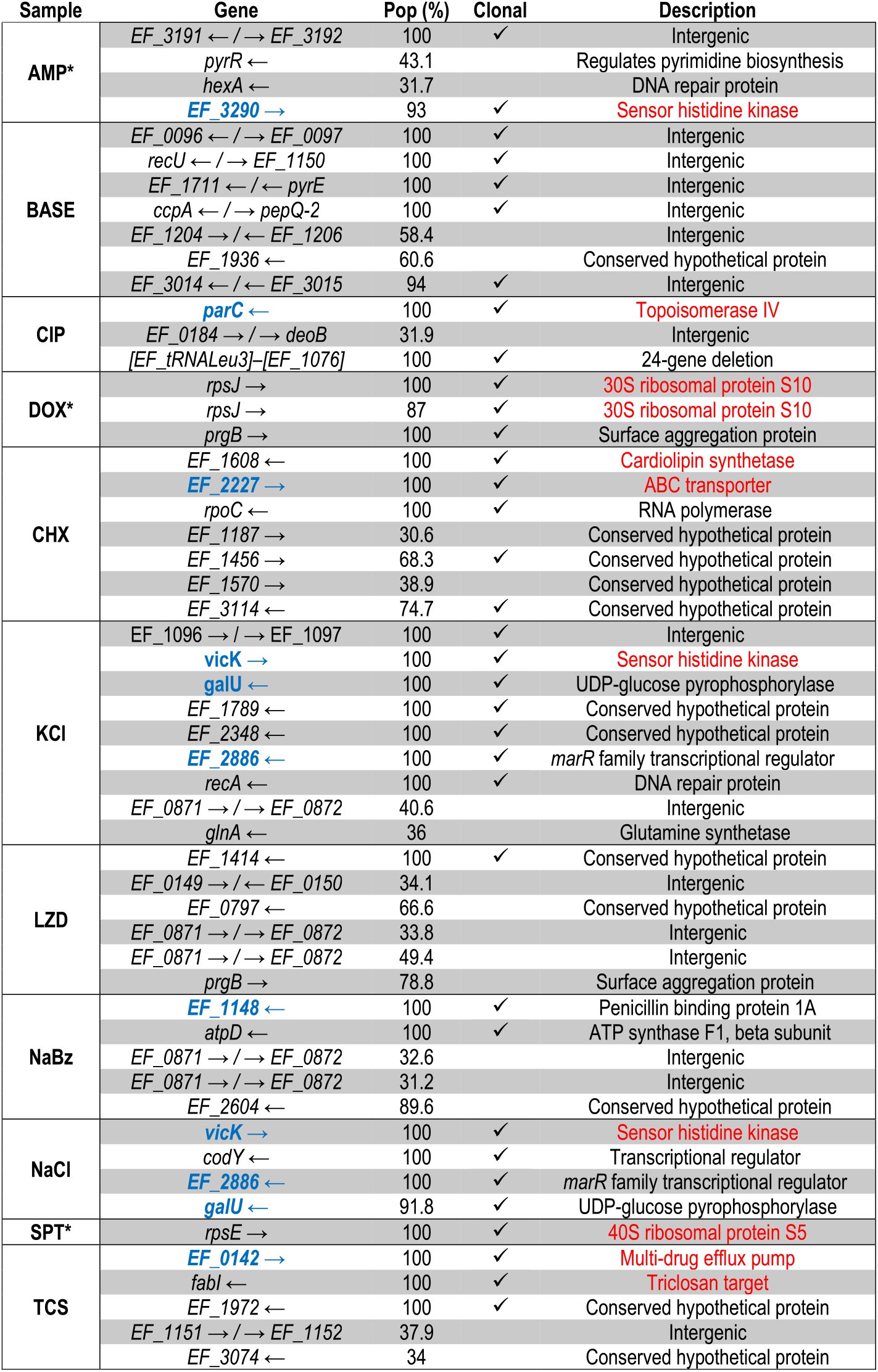
Mutations identified in selected populations. Mutations listed in red have been previously linked with resistance to the selecting condition, while genes listed in blue may confer collateral effects. Asterisk (*) identifies strains evolved and sequenced for previous work.

**TABLE 2.** Table of environmental conditions used in this study.

| Condition | Class | Mechanism of Action |
| --- | --- | --- |
| Ampicillin (AMP) | β-lactam | Inhibits cell wall synthesis |
| Doxycycline (DOX) | Tetracycline | 30S protein synthesis inhibitor |
| Spectinomycin (SPT) | Aminoglycosides | 30S protein synthesis inhibitor |
| Linezolid (LZD) | Oxazolidinone | 50S protein synthesis inhibitor |
| Ciprofloxacin (CIP) | Quinolone | DNA gyrase inhibitor |
| Nitrofurantoin (NIT) | Nitrofuran | Multiple mechanisms |
| Chlorhexidine (CHX) | Biocide | Disrupts the cell membrane |
| Triclosan (TCS) | Biocide | Inhibits fatty acid synthesis |
| Sodium Benzoate (NaBz) | Preservative | Decreases intracellular pH |
| Alkaline pH (Base) | N/A | N/A |
| Acidic pH (Acid) | N/A | N/A |
| KCl (KCl) | N/A | N/A |
| NaCl (NaCl) | N/A | N/A |

We find that isolates selected by antibiotics frequently exhibit modulated sensitivity to non-antibiotic conditions, and conversely, isolates selected by non-antibiotics often exhibit modulated sensitivity to antibiotics (Figure 1C). Sensitivity was altered in 62 percent (104/169) of condition-mutant pairs, with 58 percent (91/156) corresponding to collateral effects (i.e. modulated resistance to a stressor other than that used for selection). Collateral sensitivity is more common (58 percent, 53/91) than collateral resistance (42 percent, 38/91), though all 13 isolates exhibited both collateral resistance and collateral sensitivity to at least 2 distinct conditions.

We next asked whether the resistance profiles selected by different conditions show statistical similarities. One might hypothesize, for example, that profiles selected by chemically similar stressors would be strongly correlated with one another. On the other hand, correlations between profiles could also arise if different stressors are associated with promiscuous resistance determinants-for example, multidrug efflux pumps(46)-even for conditions that are chemically dissimilar. Indeed, we found strong (linear) correlations between the resistance profiles selected under many different pairs of conditions (Figure 2A). For example, profiles selected by NaCl are significantly correlated with those selected by acidic conditions, basic conditions, and NaBz. In addition, profiles selected by doxycycline, a protein synthesis inhibitor, are correlated with those selected by other structurally dissimilar compounds, including two antibiotics (LZD and ciprofloxacin (CIP)) as well as the antiseptic CHX. Overall, correlations between pairs of selecting conditions are dominated by positive correlations (62/78 pairs), including in all 9 pairs eclipsing the significance (*p* < 0.01) threshold. Similarly, we asked whether resistance levels between pairs of different *testing* conditions were correlated across different isolates. We found anticorrelations to be more common between testing conditions (33/78 pairs), including in two of the three pairs eclipsing significance (*p* < 0.01). Specif cally, we found negative correlations between resistance to NaCl and basic conditions and between CIP and TCS, but positive correlations between CIP and spectinomycin (SPT).

**FIG 2.**
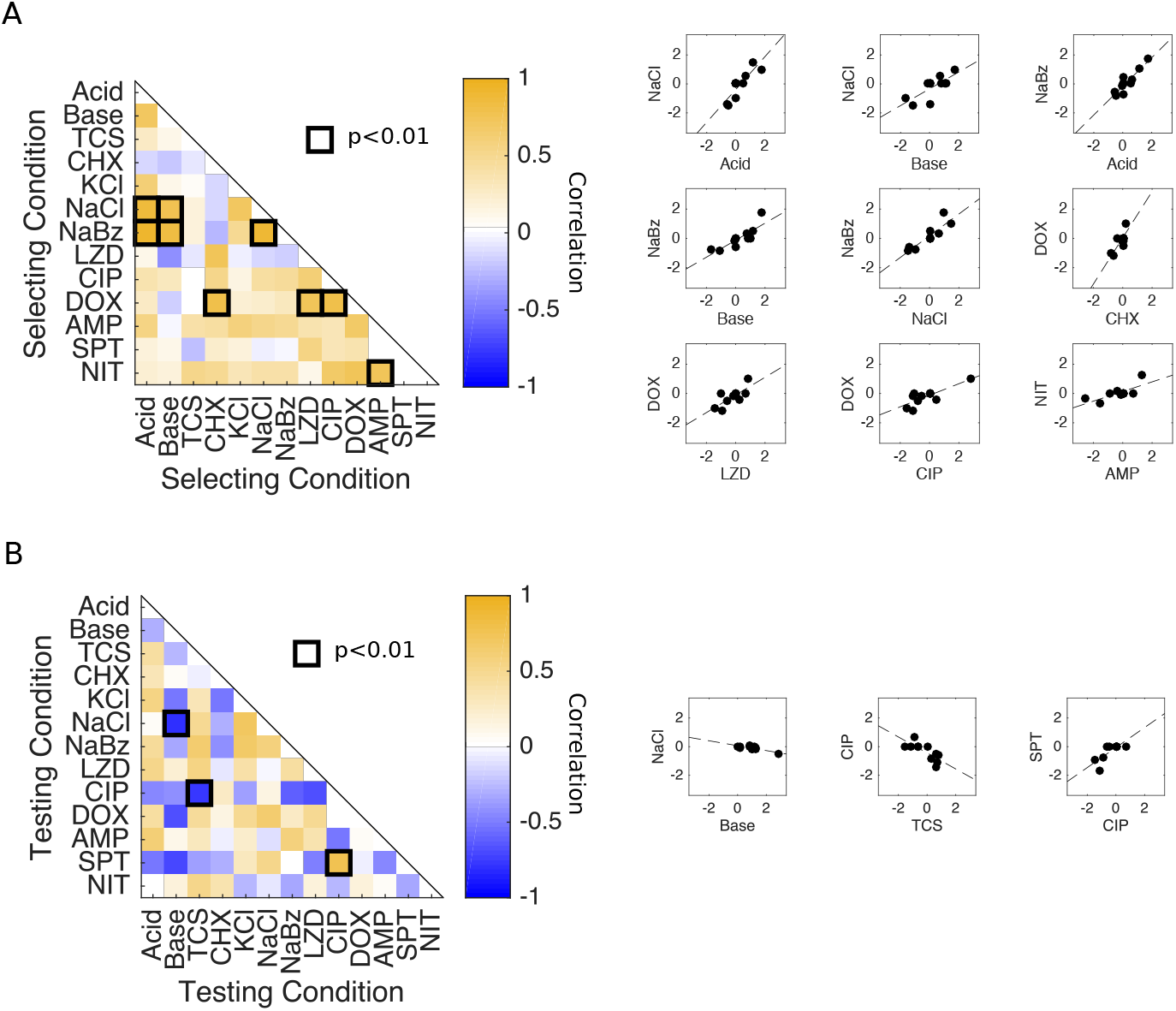
Correlations between collateral effects under different selecting or testing conditions. A. Left panel: Pearson correlation coefficient between collateral profiles (i.e. columns of the matrix in Figure 1B) selected under different conditions. Dark squares highlight significant correlations (*p* < 0.01), which are also shown as scatter plots. Right panels: pairwise scatter plots of resistance profiles selected by different conditions. Each point represents the measured resistance to a single stressor in isolates selected by the conditions on the horizontal and vertical axes. B. Left panel: Pearson correlation coefficient between resistance levels to a particular testing condition (i.e. rows of the matrix in Figure 1B) across the ensemble of isolates. Dark squares highlight significant correlations (*p* < 0.01). Right panels: pairwise scatter plots of resistance levels to different conditions. Each point represents a single isolate. To remove the effects of direct selection, which are typically larger in magnitude and may bias the correlations, the diagonal entries of the collateral sensitivity matrix (corresponding to resistance to the selecting condition) are removed prior to calculating all correlations.

### Strains selected by non-antibiotic pressures often carry mutations in genes known to confer antibiotic resistance or sensitivity

To identify candidate genes that may underlie changes in sensitivity to one or more environments, we performed whole genome sequencing on both single isolates and population samples from the evolved populations. Our goal here is not to definitively link genetic variations with specific phenotypes-a task that will require significant follow-up work. Instead, we hope to provide a roadmap of potential mutations that may underlie some of the observed collateral effects. To do so, we sequenced single isolates from each evolved population, an isolate evolved in media (BHI) only for 8 days, and also two individual isolates from the ancestral strains. In addition, we performed population sequencing on a sample from each population, including the media-selected control; we limit our analysis to variants estimated by population sequencing to occur with frequency greater than 30 percent. Note that samples from four of the antibiotic-selected populations (those selected by AMP, DOX, NIT, SPT) and the BHI-population were sequenced for a previous study (26) and their results are included here for comparison. In addition, we exclude sequencing from the acid-selected population, which was contaminated during preparation for sequencing, and exclude variants occurring in all sequenced strains. As a final control, we also confirmed a small number of mutations (in *rpsJ* in the DOX-selected isolate and in *parC* in the CIP-selected isolate) via PCR amplification and Sanger sequencing.

Overall, we observe significant agreement between population and single isolate sequencing; every mutation that occurs in the clonal sample also occurs with at least 68 percent frequency in the corresponding population sample (Table 1). Similarly, all mutations occurring at a frequency of at least 90 percent in the population samples also occurs in the clonal sequences. This consistency suggests that the phenotypic measurements, performed on single clones isolated from each population, are generally representative of the entire population, though more substantial population heterogeneity is apparent in a several cases (e.g. LZD).

This analysis reveals 54 mutations achieving at least 30 percent frequency in at least one population. We see as many as 9 mutations in a strain (KCl) and as few as zero (NIT). The control population selected by media alone showed no mutations above 30 percent frequency. 52 of the 54 mutations occurred on the choromosome, while both *prgB* mutations occured on the pTEF2 plasmid. In addition, we see several mutations present in genes known to confer resistance to the selecting drug. For example, the CHX-selected population contains two previously identified mutations responsible for CHX resistance, one in *EF_1608*, a cardiolipin synthetase, and one in EF_2227, an ABC transporter (33). The TCS isolate contains a mutation in *fabI*, a common TCS resistance gene and the drug’s target (47), as well as *EF_0142*, an efflux multi-drug resistance transporter (48). We identify a shared mutation between KCl and NaCl in *vicK*, a sensor histidine kinase known to confer resistance to environments such as osmotic stress, pH and temperature (49). In addition, the DOX-selected population contains two mutations in *rpsJ*, a gene known to confer resistance to the tetracycline class of antibiotics (50), and the CIP-selected population contains a *parC* mutation, a gene known to confer high levels of CIP resistance (51), both of which were reported previously (26).

In addition to mutations present in genes linked with resistance to the selecting condition, we also identified mutations in genes potentially responsible for modulated sensitivity to non-selecting conditions, including mutations to genes known to confer antibiotic resistance in populations selected by non-antibiotic stressors. For example, the NaCl- and KCl-selected populations harbor a mutation in *EF_2886*, a *marR* family transcriptional regulator. The *marR* system is known to regulate efflux pump activity and underlies resistance to a wide range of structurally diverse drugs (52, 53, 54). Recent work indicates that increased efflux activity comes with a trade-off, as corresponding changes in the proton-motive force can induce sensitivity to aminoglycoside antibiotics (17). Consistent with these findings, we find that isolates selected by NaCl exhibit increased sensitivity to the aminoglycoside SPT. The *marR* system is also known to confer resistance to oxidative stress, similar to TCS (55, 54); it is perhaps not surprising, then, that we observe TCS resistance in populations selected by either NaCl or KCl. We also identify a mutation in EF_1148, a penicillin binding protein (PBP), in isolates selected by NaBz. Mutations in PBPs are known to confer resistance to ?-lactam antibiotics (56), and indeed we observe cross-resistance to AMP in isolates from the NaBz-selected population.

Finally, we identify mutations in genes that have been previously linked with collateral sensitivity or resistance to antibiotics, though we observe phenotypes that differ from those expected. For example, the *marR* mutation in KCl and NaCl, the *EF_2227* mutation in CHX, and the *EF_0142* mutation in TCS are all related to efflux pumps, which are known to confer resistance to a wide array of antibiotics and biocides, especially tetracyclines and quinolones (17, 53); surprisingly, we see no increased resistance to these antibiotics in the corresponding isolates (Figure 1). Additionally, NaCl and KCl share a mutation in *galU* which is known to confer pleiotropic effects (57) and AMP resistance (58), though we see no increase in AMP resistance in isolates from the same population (Figure 1). These discrepancies could arise for several reasons. First, while we observe mutations in genes linked with drug resistance, the specific mutations are not necessarily the same. For example, the study of *EF_2227* focuses on the full gene knockout while we observe a single nonsynonymous substitution). On the other hand, the discrepancies could also be explained by epistatic effects that potentially differ in different genetic backgrounds, giving rise to variable phenotypes. It is possible that the isolate selected for phenotyping represents a rare variant of the population and therefore is not well-described by the population sequencing, though the relatively high frequencies estimated for most variants suggests this explanation is unlikely in many cases. A full list of all identified mutations is available with more details in the SI.

### Selection by chlorhexidine or triclosan frequently sensitize bacteria to at least one antibiotic

Previous studies have shown that collateral profiles may be highly variable, even when selection is performed multiple times under the same conditions (25, 26). To estimate this variability for non-antibiotic stressors, we evolved 32 replicate populations to each of two antimicrobials, TCS and CHX, for a total of 22 days (approximately 170 generations). TCS is an antimicrobial agent found in numerous consumer products, including soaps, body washes, and toothpastes. It has been linked with cross-resistance to antibiotics in multiple species (35) and was recently shown to induce resistance to antibiotics both in vitro and in vivo (59). CHX is an antimicrobial found in many disinfectants and commonly used as a general antiseptic in hospitals. CHX exposure has been linked with increased resistance to daptomycin in *E.faecium*, a closely related enterococcal species (33). Following the laboratory evolution to each condition, we measured the resistance profiles for single isolates from each population to all 13 environmental conditions (Figure 3). Surprisingly, isolates selected by each condition frequently exhibit collateral sensitivity to other agents, with 15/32 CHX isolates and 25/32 TCS isolates showing sensitivity to at least one antibiotic. In addition, all 32 CHX isolates showed strong sensitivity to TCS, while half of the 32 TCS isolates show cross-resistance to CHX.

**FIG 3.**
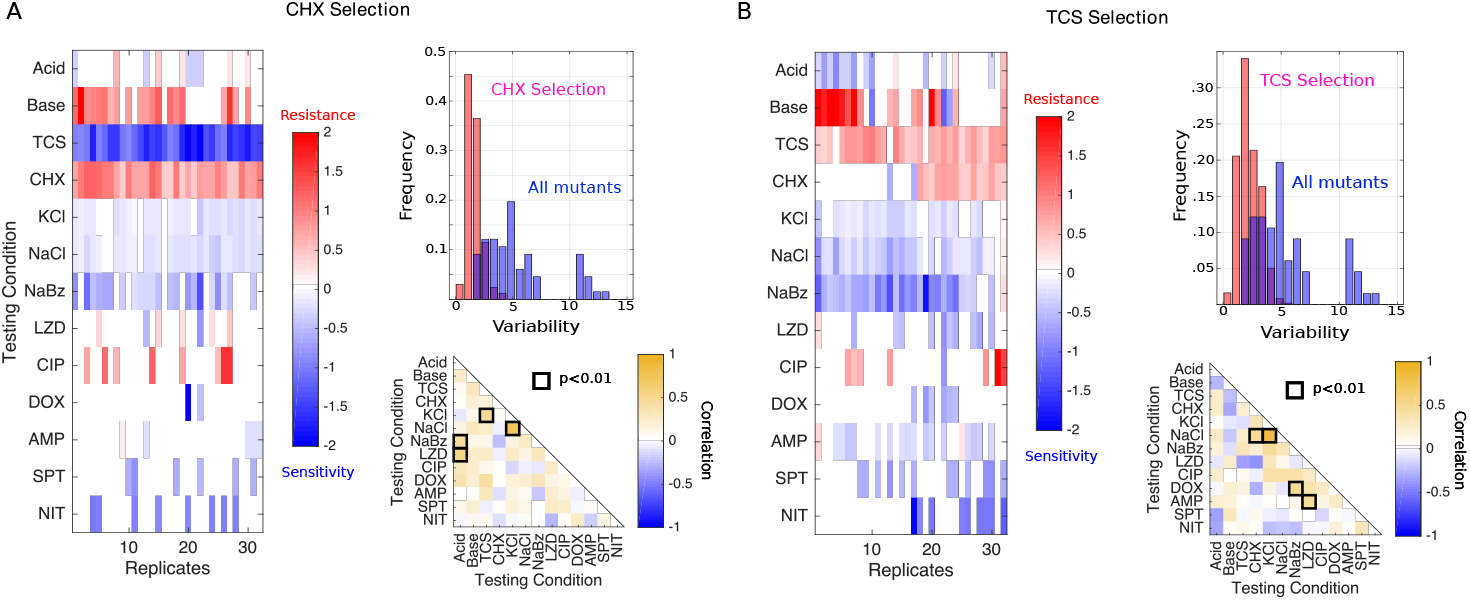
Isolates selected by CHX or TCS are variable but often exhibit increased sensitivity to antibiotics. Collateral resistance profiles for 32 independent populations evolved to either CHX (A) or TCS (B). Left panels: Resistance (red) or sensitivity (blue) to each condition (rows) is quantified using the (log_2_-transformed) fold change in the IC_50_ for the selected isolate relative to that of ancestral (V583) cells. Color scale ranges from-2 (4x decrease in IC_50_, blue) to +2 (4x increase in IC_50_, red). Top right: Histogram of variability in collateral profiles for isolates selected by CHX / TCS (red) or for the isolates in Figure 1 (spanning all conditions, blue). Variability for each collateral profile is defined as the Euclidian distance between that profile and the centroid formed by the relevant ensemble of profiles. Bottom right: Pearson correlation coefficient between resistance levels to a particular testing condition across the ensemble of isolates.

To quantify variation within an ensemble of collateral profiles, we considered each profile as a 13-dimensional vector, with each component representing resistance to a particular environmental condition. To estimate variability within the ensemble, we calculated the mean pairwise (Euclidean) distance, 〈*d_p_*〉, across all pairs of profiles in the ensemble. While collateral profiles of isolates selected by TCS (〈*d_p_*〉 = 2.2) and CHX (〈*d_p_*〉 = 1.6) both exhibit isolate-to-isolate variability, it is considerably smaller than the variability observed across all conditions (〈*d_p_*〉 = 5.2). In addition, the distribution of pairwise distances between isolates selected by the same condition (TCS or CHX) is considerably more narrow that the distribution across all isolates (Figure 3, upper right insets). We also tested for correlations between resistance levels to pairs of stressors across the ensemble of isolates for each condition. Not surprisingly, the correlations between pairs of stressors vary substantially depending on the selecting conditions used to generate the isolates (compare insets in Figure 3A, 3B). For example, resistance to KCl is correlated with resistance to TCS following CHX selection (Figure 3A, lower right) but weakly anticorrelated in TCS-selected isolates (Figure 3B, lower right). On the other hand, there are rare pairs of environments-such as NaCl and KCl-where resistance is strongly correlated in all sets of isolates, likely reflecting the extreme chemical similarity between the stressors.

### Sequential rounds of antibiotic and non-antibiotic selection can promote sensitivity

Our results indicate that both collateral sensitivity and cross resistance are surprisingly common in the evolved lineages. Selection by one condition (by definition) leads to resistance to that condition, but it frequently sensitizes the population to multiple other conditions. In fact, our experiments showed that selection by one stressor led to increased sensitivity to between 3 and 7 other conditions (Figure 1B). Unfortunately, these increased sensitivities are also accompanied by frequent cross-resistance, placing limits on the number of sensitivities that can be selected by any one condition.

However, we hypothesized that it might be possible to circumvent those limitations by using a sequence of two stressors. While this sequential selection is likely to produce resistance to, at minimum, the two selecting conditions, it’s possible that judiciously chosen conditions could lead to more sensitivities than either condition alone-in effect harnessing the orthogonal sensitizing effects of particular pairs of selective forces. To guide our search, we first calculated the expected number of sensitivities following sequential selection by each pair of conditions under the naive assumption that phenotypic effects are purely additive. Because resistance is measured on a log scale, the assumption of additivity means that relative changes in IC_50_ (or similar) are multiplicative; for example, if conditions 1 and 2 each reduce IC_50_ to 40 percent of the value in ancestral cells, their sequential application would reduce IC_50_ to 16 percent. We note that such null models are imperfect, as they fail to capture epistasis and known hysteresis in evolutionary trajectories (see, for example, (23)). Here we use the null model only to identify candidate condition pairs for further experimental investigation. Under these additivity assumptions, the number of sensitivities is expected to increase for most pairs of stressors; that is, assuming additivity of the measured sensitivity profiles, sequential exposure to pairs of stressors is often predicted to sensitize the population to more stressors than exposure to either single agent alone (Figure 4A). In three cases (LZD-NaCl, LZD-NaBz, and NIT-SPT), the number of sensitivities is expected to increase by three or more, providing a substantial benefit over the single agent selecting conditions.

**FIG 4.**
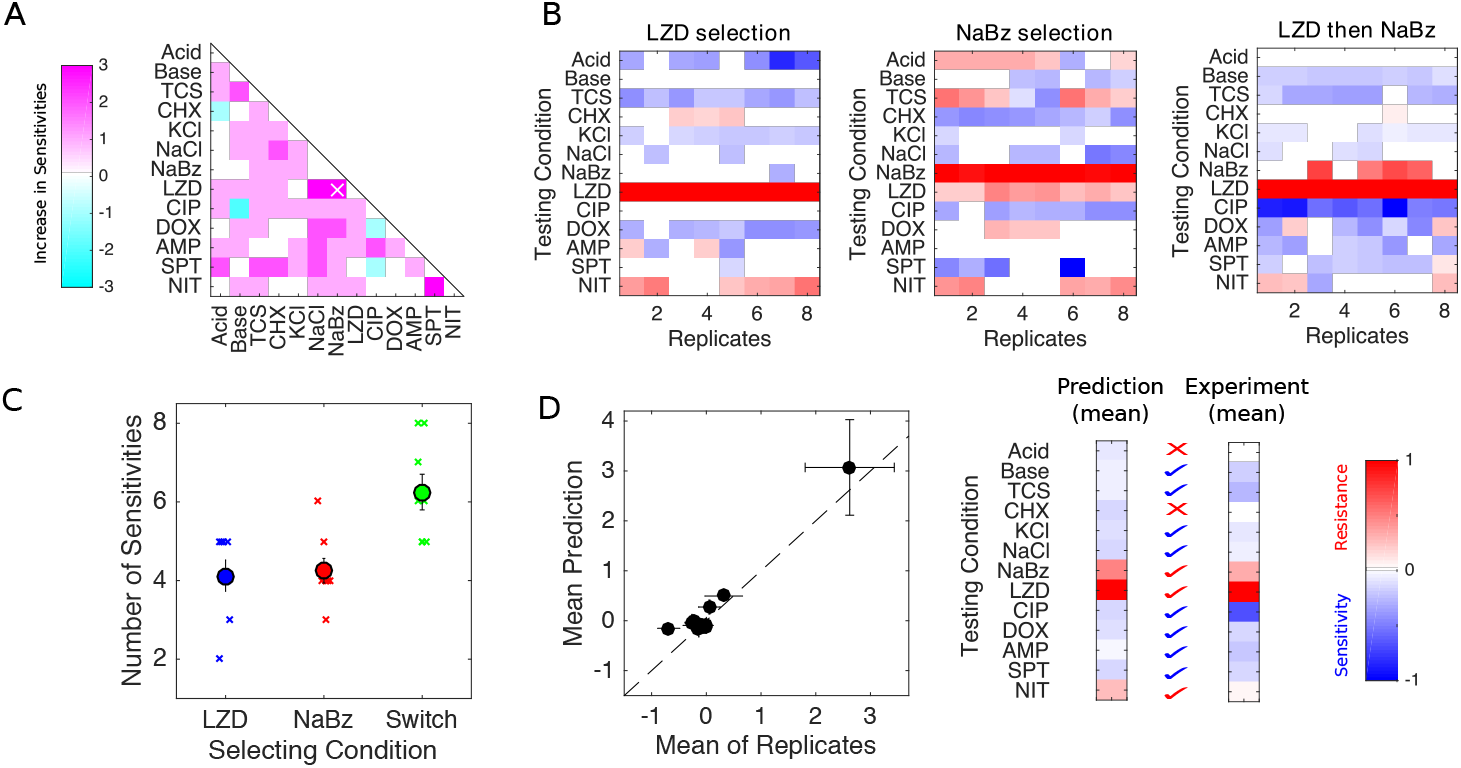
Selection in alternating environments can induce sensitivity to more stressors than selection in single environments. A. Predicted change in number of sensitivities following sequential evolution using pairs of conditions. Change is positive if sequential evolution is predicted to result in more sensitivities than evolution in either component alone. Predictions assume additivity of (log-scaled) resistance profiles. A sequence of LZD and NaBz (white x) is predicted to give the maximum increase in sensitivities. B. Resistance profiles for replicate evolution experiments (8 per condition) in LZD only, NaBz only, or a two-step sequence consisting of LZD selection followed by NaBz selection. Resistance (red) or sensitivity (blue) to each condition (rows) is quantified using the (log_2_-transformed) fold change in the half-maximal inhibitory concentration (IC_50_) for the selected isolate relative to that of ancestral (V583) cells. C. Left panel: Isolates evolved in the alternating environment (“switch” between LZD then NaBz, green) exhibit sensitivity to more environments than isolates selected in each environment alone (blue and red; *p* < 0.01, Wilcoxon rank sum test for pairwise comparisons between LZD and switch and between NaBz and switch. D. Left: scatter plot comparing the mean collateral profile of isolates from the alternating selection (“experiment”) and mean collateral profiles predicted by an average (linear sum) of profiles generated in the single environments (“prediction”). Right panel: heat maps (color scale same as for panel A). Check marks indicate correctly predicted sensitivity (blue) or resistance (red). X indicates incorrect prediction.

To test these predictions experimentally, we focused on the pair LZD, a protein synthesis inhibitor, and NaBz, a commonly used food preservative. Our original selection experiments showed that selection in LZD led to 5 sensitivities and NaBz led to 4 sensitivities; the sensitivities are largely non-overlapping, and sequential selection is therefore predicted to an increase in the number of sensitivities. To test this prediction, we performed experimental evolution on eight replicate populations to each of 3 conditions: LZD alone, NaBz alone, and a two-phase sequence consisting of LZD evolution followed by NaBz evolution. For convenience, we limited each evolution phase to 10 days (70-80 generations), making this considerably shorter than the original adaption in Figure 1. We then tested an isolate from each population for modulated resistance to each of the 13 environmental conditions (Figure 4B).

The isolates selected by LZD or NaBz alone had sensitivity profiles that are similar, but not identical, to those selected in the original experiment (Figure 1). For both conditions, the single agent evolution led to increased sensitivity to an average of approximately 4 conditions (Figure 4C). Strikingly, however, evolution in the LZD-NaBz sequence (“switch”) sensitized the isolates to more than 6 conditions on average, with some isolates exhibiting sensitivity to eight conditions.

To test the quantitative accuracy of the null model, we generated an ensemble of plausible resistance profiles for the sequential selection experiment. Each profile in the predicted ensemble corresponds to the mean of one pair of profiles, with one member of the pair drawn from the LZD only selection (Figure 4B, left) and one drawn from the NaBz only selection (Figure 4B, middle). The mean pro1le in this ensemble agrees surprisingly well with the mean pro1le measured in the LZD-NaBz evolution (Figure 4D).

## DISCUSSION

These results provide a systems-level picture of the phenotypic trade-offs accompanying evolved resistance to antibiotic and non-antibiotic stressors in an opportunistic pathogen. We find that collateral resistance and collateral sensitivity are surprisingly pervasive across conditions, underscoring the need to better understand how adaptation to non-antibiotic environments may contribute to drug resistance. These widespread collateral effects raise the question of whether frequently encountered stressors-food additives, preservatives, biocides, or simply common elements of natural environments-may steer bacteria toward multidrug resistance, and in turn, whether there may be an unappreciated role for these agents in slowing or reversing resistance. As proof-of-principle, we showed experimentally that sequential adaptation to different environments can be used to sensitize bacterial to antibiotics, a consequence of the largely non-overlapping sensitivities induced by each agent alone.

The goal of this study was to investigate patterns of resistance between antibiotics and non-drug stressors at a phenotypic level. By taking a systems-level view, we hoped to gauge the prevalence collateral sensitivity and assess the potential of non-antibiotic agents for evolutionary steering. This approach comes with obvious drawbacks, and it indeed leaves us with many unanswered questions. Most notably, it is vitally important to understand the molecular and genetic mechanisms facilitating these overlapping resistance profiles, though doing so on a broad scale is not easily done in one study. There are many well-known examples of molecular mechanisms that confer nonspecific resistance to structurally unrelated compounds in bacteria, including a number of multidrug resistance transporters and efflux pumps (52, 48, 60, 46). On the other hand, collateral sensitivity in bacteria remains much less understood, even between antibiotics. Recent evidence suggests these sensitivities may be governed by target mutations that induce global changes in gene regulation or by mutations altering drug uptake and efflux (61). Similar mutations appear in our evolved mutations, suggesting that these mechanisms may also underlie many of the observed collateral effects between antibiotic and non-antibiotic stressors. However, definitely linking particular mutations with phenotypic effects will require considerable follow-up work to disentangle, for example, the potential effects of mutational epistasis and genetic background on drug resistance phenotypes.

We have shown experimentally that sequential adaptation to antibiotic and nonantibiotic conditions can sensitize bacteria to more environments than either agent alone. While we focus here on a clinically relevant bacterial species, it is not clear the these results will generalize to other species. We used a simple additive model to identify candidate environmental pairs for sequential selection. While the model gave surprisingly accurate predictions in these experiments, it will clearly fail when effects of epistasis or evolutionary hysteresis are strong. On the other hand, if epistasis effects are approximately symmetric about zero or typically small relative the core effects of additivity, similar null models may still prove useful for finding environmental pairs that increase the number of sensitivities, though the predictions of specific profiles are likely to become increasingly inaccurate. Long-term application will therefore require continued experimental mapping of the collateral sensitivity profiles selected by increasingly complex and realistic environmental conditions.

## MATERIALS AND METHODS

### Strains, antibiotics, non-antibiotics and media

All mutants were derived from *E. faecalis* V583, a fully sequenced vancomycin-resistant clinical isolate (62). The 13 conditions used to select mutants are listed in Table 1. Antibiotics were prepared from powder stock and stored at −20C with the exception of ampicillin, which was stored at −80C. TCS, CHX and NaBz were prepared from powder stock and stored at −20C. Acid (pH≈1.5) and Base (pH> 10.5) stock solutions were prepared by titrating HCl and NaOH, respectively, into BHI medium. These stock solutions were mixed in appropriate volumes with standard BHI (pH≈7) to create selecting media for evolution experiments. Saturated KCl and NaCl stock solutions were prepared by dissolving KCl and NaCl into BHI medium. As with acid and base, appropriate mixtures of saturated KCl and NaCl solutions were mixed with standard BHI medium. Evolution and IC_50_ measurements were conducted in BHI medium alone with the exception of daptomycin, which requires an addition of 50 mg/L calcium for antimicrobial activity.

### Laboratory Evolution Experiments

Evolution experiments were performed in 96-well plates with a maximum volume of 2 mL and a working volume of 1 mL BHI. Each day, at least three replicate populations were each grown in a different concentrations of the selecting agent. The concentrations were chosen to include both sub- and super-inhibitory concentrations. After 20-23 hours of incubation at 37C, aliquots (5 *μ*L) from the population that survived (OD>0.3) the highest concentration were added to a new series of wells and the procedure was repeated for 50-60 days (350-450 generations). Note that isolates from antibiotic selection experiments (see (26)) were evolved for only 8 days, in part because resistance to antibiotics increased much more rapidly than resistance to other agents, such as NaCl. We chose longer timescales for the non-antibiotic conditions to ensure resistance to the selecting condition increased by approximately 2x or more in each case. On the final day of selection, we plated a sample from each population on BHI agar plates, isolated a single colony from each plate, and stored the remaining population volume at −80C in 30 percent glycerol.

### Measuring Drug Resistance and Sensitivity

IC_50_ measurements for each condition/drug were performed in triplicate for each isolate (except in the case of the ancestral wild-type strain, which was performed in replicates of 8) in 96-well plates by exposing mutants in different wells to 6-10 concentrations of drug, typically in a linear dilution series prepared in BHI medium. After 12 hours of growth at 37C, the optical density at 600 nm (OD) was measured using an Enspire Multimodal Plate Reader (Perkin Elmer) with an automated 20-plate stacker assembly.

Each OD reading was normalized to by the OD reading for the same isolate in the absence of drug. To quantify resistance, the resulting dose response curve was fit to a Hill-like function *f*(*x*) = (1 + (*x/K*)^*h*^)^−1^ using nonlinear least squares fitting, where K is the half-maximal inhibitory concentration (IC_50_) and h is a Hill coefficient describing the steepness of the dose-response relationship. A mutant strain was deemed collaterally sensitive (resistant) if its IC_50_ varied by more than 3*σ_WT_* from that of the ancestral strain, where *σ_WT_* is the uncertainty (standard error across 8 replicates) of the IC_50_ measured in the ancestral strain. Note that all estimates of IC_50_ in the ancestral (“wild-type”) strain, across all replicates and for all conditions, are contained in this ±3*σ_wT_* range, which suggests that there are unlikely to be false positives in designating isolates as sensitive or resistant.

## Supporting information

SI: Sequencing_VariantCalls

## ACKNOWLEDGMENTS

This work is supported, in part, by the National Science Foundation (NSF No. 1553028 to KW), the National Institutes of Health (NIH No. 1R35GM124875-01 to KW), and The Hartwell Foundation for Biomedical Research (to KW). The format for this preprint is adapted from the American Society for Microbiology (ASM) template available on Overleaf.com.

